# spatialGE: A user-friendly web application to democratize spatial transcriptomics analysis

**DOI:** 10.1101/2024.06.27.601050

**Authors:** Oscar E. Ospina, Roberto Manjarres-Betancur, Guillermo Gonzalez-Calderon, Alex C. Soupir, Inna Smalley, Kenneth Tsai, Joseph Markowitz, Ethan Vallebuona, Anders Berglund, Steven Eschrich, Xiaoqing Yu, Brooke L. Fridley

## Abstract

Spatial transcriptomics (ST) is a powerful tool for understanding tissue biology and disease mechanisms. However, its potential is often underutilized due to the advanced data analysis and programming skills required. To address this, we present spatialGE, a web application that simplifies the analysis of ST data. The application spatialGE provides a user-friendly interface that guides users without programming expertise through various analysis pipelines, including quality control, normalization, domain detection, phenotyping, and multiple spatial analyses. It also enables comparative analysis among samples and supports various ST technologies. We demonstrate the utility of spatialGE through its application in studying the tumor microenvironment of melanoma brain metastasis and Merkel cell carcinoma. Our results highlight the ability of spatialGE to identify spatial gene expression patterns and enrichments, providing valuable insights into the tumor microenvironment and its utility in democratizing ST data analysis for the wider scientific community.

## Background

The ability to quantify spatially-resolved gene expression within tissues has been enhanced with the development and commercialization of various spatial transcriptomics (ST) technologies^1, 2^. Increasingly, researchers are including spatial transcriptomics data as an additional modality to complement the power of single-cell transcriptomics and localize cell types or tissue niches of interest within the larger context of the tissue architecture^2, 3^. In other cases, ST has been used independently as a hypothesis-generating assay for later exploration of cell-cell interactions via functional “wet-lab” experiments^4^. The various ST platforms are available at different levels of molecular resolution (i.e., curated gene panels vs. whole transcriptome) and cellular resolution (10-100s cells per spot/ROI to single cell-level/subcellular quantification) ^1^, allowing data acquisition for a wide range of questions from large scale, tissue niche expression patterns to cell-cell communication^5^. The popularity of ST among the scientific community (**Fig. 1**) has also prompted the development of a myriad of algorithms, including bioinformatic pipelines to facilitate data quality control, cell type discovery, tissue domain detection, cell phenotyping, and cell-cell communication^6, 7^.

**Figure 1.**
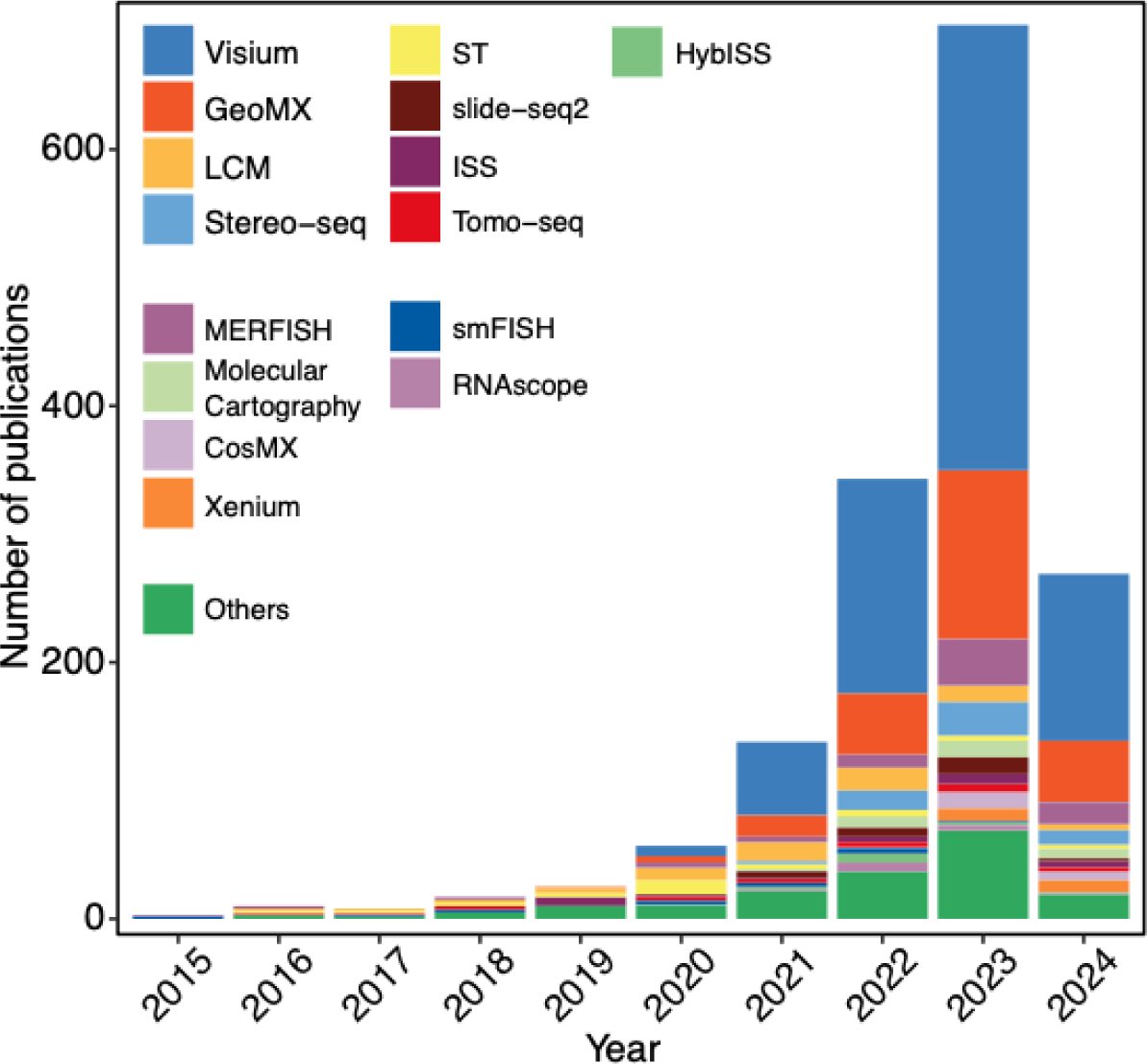
The number of publications per year involving spatial transcriptomics data. Publication records extracted from The Museum of Spatial Transcriptomics^2^ (June 2024).

Despite the enormous availability of analytical methods, expertise in analyzing high-dimensional data and programming is required for their utilization, limiting the initial exploratory data analysis for scientists. Additionally, expertise in spatial data analysis is often necessary for advanced bioinformatic and statistical analysis of ST data. As a result, most tissue biologists depend on bioinformaticians for basic and advanced analysis. Similarly, bioinformaticians need to run many analyses iteratively with input from biologists to explore potential hypothesis-testing directions. Hence, user-friendly software is necessary for non-data scientists to explore and leverage their ST data to the fullest. Efforts to automatize routine analysis in a user-friendly manner have been provided by many companies commercializing ST platforms; however, these solutions are often proprietary and severely limited in scope. Open-source solutions are also available with varying degrees of user-friendliness, including applications launched from the command line^8^ or integrated into cloud services^9^ to ready-to-use web applications^10, 11^. The current web server-based applications feature either a fixed workflow^11^ or an interface to build pipelines from a combination of available tools^10^. While these server-based applications are an important improvement towards making ST data analysis accessible to researchers, there is a need for ST analysis workflows with improved accessibility and guidance to navigate the complexities of spatial analysis. Furthermore, comparative analysis is also limited in current server-based solutions, which generally allow analysis of a single sample at a time.

With all these challenges in mind, we have developed a web application that wraps the functionality of the spatialGE^12^ R package, as well as other analytical tools, to provide a comprehensive, user-friendly, point-and-click platform for the analysis of spatial transcriptomics. We refer to this web application as spatialGE. The spatialGE application is modular-based and carefully guides the user through various analysis pipelines, including quality control, normalization, domain detection, phenotyping, and multiple spatial analysis. spatialGE also facilitates comparative analysis among samples, including qualitative integration with sample-level metadata and the visualization of results in the context of the tissue images. Lastly, spatialGE can be used across various ST technologies, whereby only a table with gene counts for all cells/spots, and a table with the spatial coordinates are required. In addition to accepting the generic formatted files, spatialGE currently takes output generated from 10X Genomics’ Visium and NanoString’s CosMx SMI platforms.

## Results

The application spatialGE guides users through an ST analysis pipeline using a modular format (**Fig. 2**). This allows the users to conduct specific parts of the spatialGE pipeline and assess the results of each module separately. The modules in the web application include data entry (*Import data*), filtering and normalization (*QC and data transformation*), gene expression plotting (*Visualization*), spatial heterogeneity analysis (*Spatial heterogeneity*), spatial-based gene set analysis (*Spatial gene set enrichment*), spot/cell clustering (*Spatial domain detection*), gene marker detection (*Differential expression*), and gene expression spatial gradient analysis (*Spatial gradients*). To help guide the users, the modules become available in the interface once certain analysis requirements are met. For example, the *QC and data transformation* module becomes available only after data has been imported, and the *Visualization* module is enabled only after data has been transformed. Comprehensive documentation is available within each module, including user manuals and “info-tags” that provide details about the function of each control in the user interface.

**Figure 2.**
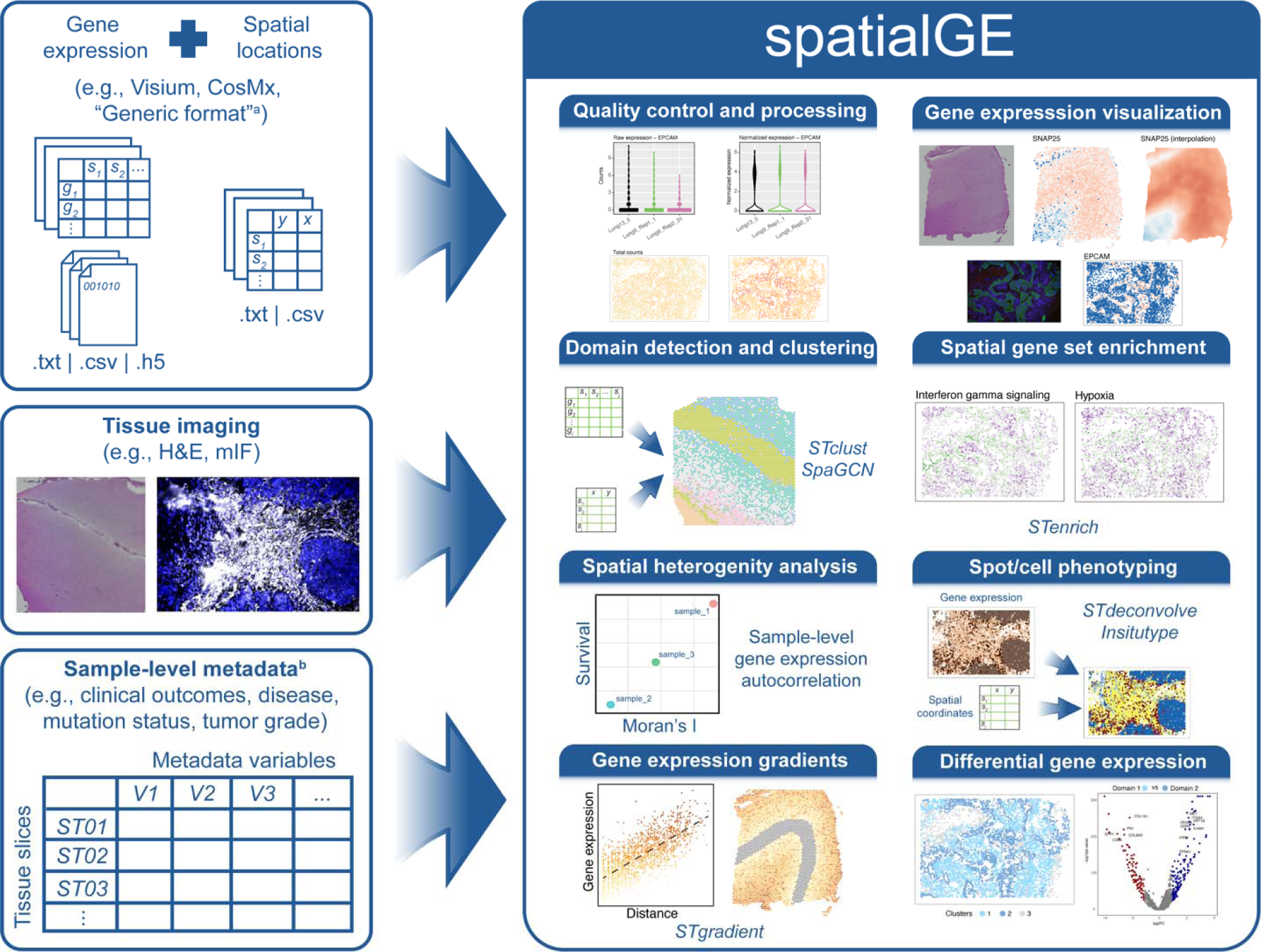
The functionality of spatialGE. The web application can take ST data directly from the Space Ranger tool for Visium (.h5) or CosMx. The “generic format” is a third input option, accepting .txt or .csv tables with gene expression and spatial coordinates. Tissue images can also be uploaded. Sample-level metadata provided by the user can be easily input in spatialGE, facilitating comparative analysis. Once the ST data has been loaded, spatialGE guides the user through the different steps of an exploratory analysis pipeline, including quality control (gene and spot/cell filtering), count normalization, clustering, spatial analysis, and visualization.

Aside from its analytical capabilities, users set up an account that can house multiple projects. For each project, users can upload spatial transcriptomics data from multiple samples, and associated analyses are saved within the project within the user’s account (see *Methods– Software architecture*). Analysis in spatialGE can be conducted in the background with the functionality to notify users when an analysis has been completed via e-mail. Results and parameters used in each module can be downloaded to facilitate reproducibility.

### spatialGE modules

#### Import data module

Within the user’s account, the user begins by creating a project. The project holds the imported ST data and results derived from the analysis modules. Currently, spatialGE takes data in three formats: Generic, Visium, and CosMx. A sample (i.e., tissue slice) with ST data in generic format consists of an untransformed gene expression count matrix (.csv or .tsv) and a spatial coordinates file (.csv or .tsv). The gene expression matrix contains genes in rows and spots/cells in columns, with the first column of the matrix containing the gene names (non-duplicated HUGO symbols). The spatial coordinates file has three columns. The first column contains the spot/cell IDs, matching the column names in the expression matrix. The second and third columns include the Y and X values indicating the spatial coordinates of the spots/cells. Users can also upload an accompanying tissue image that can be displayed next to the gene expression plots.

The Visium format consists of files output by the 10X Space Ranger pipeline. For each sample (i.e., a single Visium slide area), the user provides the .h5 gene expression file and the coordinates file (“tissue_positions_list.csv”). Optionally, users can upload the “tissue_hires_image.png” and “scalefactors_json.json” files to display the tissue image next to expression plots. The CosMx format takes gene expression (“exprMat_file.csv”) and coordinates (“metadata_file.csv”) files that contain data for each slide, with a slide containing several fields of vision (FOVs). Internally, the spatialGE splits the data into the FOVs, with analyses completed at the FOV level.

An important feature of spatialGE is the ability to make comparative analyses across several samples and display metadata associated with each sample (e.g., *Spatial heterogeneity* module). Sample-level metadata is provided as a .csv or .xlsx file with one row per sample. The first column contains sample names, and subsequent columns contain the variables. Alternatively, users can optionally enter the metadata directly within the *Import data* module. Examples of metadata include overall survival, tissue type, tissue preservation method, or any other attribute of the sample.

#### QC and data transformation module

After completing the data entry (*Import data* module), the *QC and data transformation* module is enabled. The module provides a summary table of the number of genes and counts detected at the spots/cells. Values in the summary are updated after filtering and quality control have been performed in the corresponding section (*Filter data* tab). In the *Filter data* tab, users can remove entire samples from the project or spots/cells based on gene counts. Functionality is available to remove all counts for specific genes, such as mitochondrial, ribosomal genes, or negative control probes. Advanced users can filter out genes by typing regular expressions in the graphical interface.

The two normalization methods in the R package are also available in spatialGE (*Normalize data* tab). The first is a library-size normalization followed by log-transformation (log 1+counts). The second is the SCTransform method^13^, a widely used normalization in single-cell studies. The interface generates visualizations to assess the effect of the selected normalization method, informing the user to take appropriate action. Principal component analysis of “pseudobulk” ST samples (i.e., gene-level sum of counts for all spots/cells) is available, as well as visualizations of total counts in the spatial context of the sample (*Quilt plot*).

#### Visualization module

Gene expression at each spot/cell for one or more genes is available in the *Visualization* module. We have implemented two options introduced in the first version of the R package spatialGE^12^: Quilt plots and expression surfaces (**Fig. 2**). The quilt plot shows the raw or normalized expression of each user-selected gene at each spot/cell. The expression surface is a spatial interpolation plot generated via kriging^14^. The expression surface plot is particularly useful in technologies such as Visium, where spots are separated by 100µm.

#### Spatial heterogeneity module

Spatial heterogeneity statistics (e.g., Moran’s I, Geary’s C) can be calculated for each sample in the *Spatial heterogeneity* module. Details about the statistical framework and interpretation of metrics can be obtained from the publication describing the R package spatialGE^12^. The spatial statistics measure the tendency in the expression of a gene to be concentrated (“hotspot”) or dispersed (“uniform”) across the tissue. Once spatial statistics have been calculated for one or more genes, the values for Moran’s I and/or Geary’s C can be shown in a plot to facilitate comparative analysis. If the user has provided sample-level metadata, samples can be grouped using this information.

#### Spatial gene set enrichment module

The *Spatial gene set enrichment* module implements STenrich, a method in the spatialGE R package. The *Spatial gene set enrichment* module tests for the existence of spatial patterns (e.g., “hotspots”) at the gene set or pathway level for each sample. The STenrich algorithm provides a p-value for the statistical significance of spatial patterns across a collection of gene sets using a permutation-based approach (see *Methods–spatialGE analysis modules* for algorithm details). Currently, KEGG, GO, HALLMARK, and REACTOME gene sets are available for testing.

#### Spatial domain detection module

In spatialGE, we have implemented clustering approaches to facilitate the detection of tissue domains or niches. Currently, two spatially-informed methods are available. The STclust^12^ is a method within the spatialGE R package and allows the application of spatial weights, which moderates the gene expression differences and results in domains that are spatially continuous. We have also incorporated the SpaGCN^15^ algorithm, which uses graph convolutional networks to integrate spatial distances with gene expression. The *Spatial domain detection* module generates visualizations of the detected domains and allows visual comparative analysis across samples and each sample with its tissue image if available. In addition, if the user has detected domains with SpaGCN, the module has built-in support for detecting spatially variable genes (SVGs).

#### Differential expression module

spatialGE supports non-spatial and spatially aware tests for determining differentially expressed genes among tissue domains detected in the *Spatial domain detection* module. Traditional tests such as Wilcoxon’s rank and t-test are available (i.e., non-spatially aware methods). We have also implemented the use of mixed models with spatial covariance structures to account for the spatial correlation of gene expression measurements among adjacent spots or cells. This spatially-aware testing approach to differential gene expression analysis, called STdiff^16^, reduces the overrepresentation of small (“statistically significant”) p-values often observed when non-spatial tests are conducted on ST data.

#### Spatial gradients module

Once tissue domains have been detected (*Spatial domain detection* module), users can test for genes that show a spatial gradient of expression. A gene showing a spatial gradient exhibits higher expression in spots/cells closer to a tissue domain and lower expression in spots/cells farther from the same domain. The module implements the STgradient function from the spatialGE R package. The STgradient method is especially useful for assessing spatial gene expression patterns at the interface of tissue domains, with the advantage that STgradient does not require a priori specification of an “interface” niche (see *Methods– spatialGE analysis modules* for algorithm details).

#### Phenotyping module

The *Phenotyping* module implements two pipelines for the phenotyping of cells or spots in spatial transcriptomics experiments. The first approach, STdeconvolve^17^, allows the identification of cell types within each spot of a Visium sample. The module guides the user through model fitting to find the most likely number of latent topics (i.e., number of cell types or niches), followed by gene set enrichment analysis (GSEA) to assign biological identities to the latent topics. The GSEA uses a list of markers for each cell type, which we have derived from CellMarker 2.0^18^ and incorporated into spatialGE, avoiding needing a scRNA reference. The set list of markers will be periodically updated. However, the current lists are available to the reader in the **Additional file 1**. In the future, spatialGE will allow users to upload customized cell-type marker lists. For single-cell level technologies, such as CosMx, we have implemented Insitutype^19^. The Insitutype algorithm identifies cell types using a Bayes classifier and gene expression signatures from the cell types potentially present in the tissue of interest. spatialGE provides several built-in gene expression signatures derived from the SpatialDecon R package^20^.

### Application of spatialGE to studying the tumor microenvironment

The application spatialGE includes many modules for comprehensive exploratory analysis of data from ST technologies. To illustrate the utility of spatialGE we have focused on applying two advanced methods within the spatialGE suite: *Spatial gradients* and *Spatial gene set enrichment* for discovering gene-level and gene set-level spatial patterns. We completed data filtering, normalization (*QC and data transformation* module), tissue domain detection (*spatial domain detection* module), phenotyping, visualization, and spatial analysis (*spatial gradients* and *spatial gene set enrichment* modules) for two cancer studies. The first study contains data on melanoma brain and extra-cranial metastases tissues profiled with 10X Genomics’ Visium platform, while the second study involves a Merkel cell carcinoma study profiled with NanoString’s CosMx SMI technology.

#### Melanoma metastases

The brain is a common metastasis site in melanoma patients, often with a very poor prognosis. Advances in targeted therapy, immunotherapy, and stereotactic radiosurgery have led to improved prognoses. However, median survival is 1-2 years^21^. We used spatialGE to explore the spatial gene expression in three melanoma brain metastases and perform comparative analysis with four melanoma extra-cranial metastases samples. ST data was generated using 10X Genomics’ Visium platform. After filtering, each sample contained an average of 1333 to 6460 counts per spot and 832 to 2938 detected genes per spot (**Fig. 3 A, B**). Tissue domains were predicted for all samples using the *Spatial domain detection* module with STclust^12^. Following the identification of the tissue domains, we assigned domains as “tumor” based on the differential expression results (see Methods–*Melanoma metastasis study*) and comparison of detected domains to the H&E image. Next, we used the *spatial gradients* module to determine genes in which the expression level changes as a function of distance from the “tumor” tissue domains (**Fig. 3C**). The genes showing a significant correlation of expression with distance from tumor areas (Spearman’s rho p-value) varied from sample to sample, reflecting the unique tissue architecture of each sample (**Fig. 3D**). After completing the first analysis with the *Spatial gradients* module, one brain metastasis (sample9_LMM_F) was removed from the comparisons, given the low overlap of genes showing significant spatial gradients compared to the rest of the samples in the study. We observed that in the two brain metastases left in the analysis, a negative Spearman coefficient for the expression of *S100A1* was present, indicating higher expression as the distance from the tumor domain decreased (**Fig 3E**). Analysis with the *Differential expression* module indicated that the genes *CHI3L1*and *CHI3L2* were upregulated in the astrocyte-dominated regions of sample1_MB14_A. The presence of astrocytes in this region was also observed using gene expression deconvolution (**Additional file 2**) in the *Phenotyping* module (STdeconvolve^17^). We tested for expression gradients with respect to astrocyte-dominant areas in sample1_MB14_A, which revealed higher expression of the chemokine ligand *CCL18* in areas far from the tumor domains (**Fig 3F**; **Additional file 3**).

**Figure 3.**
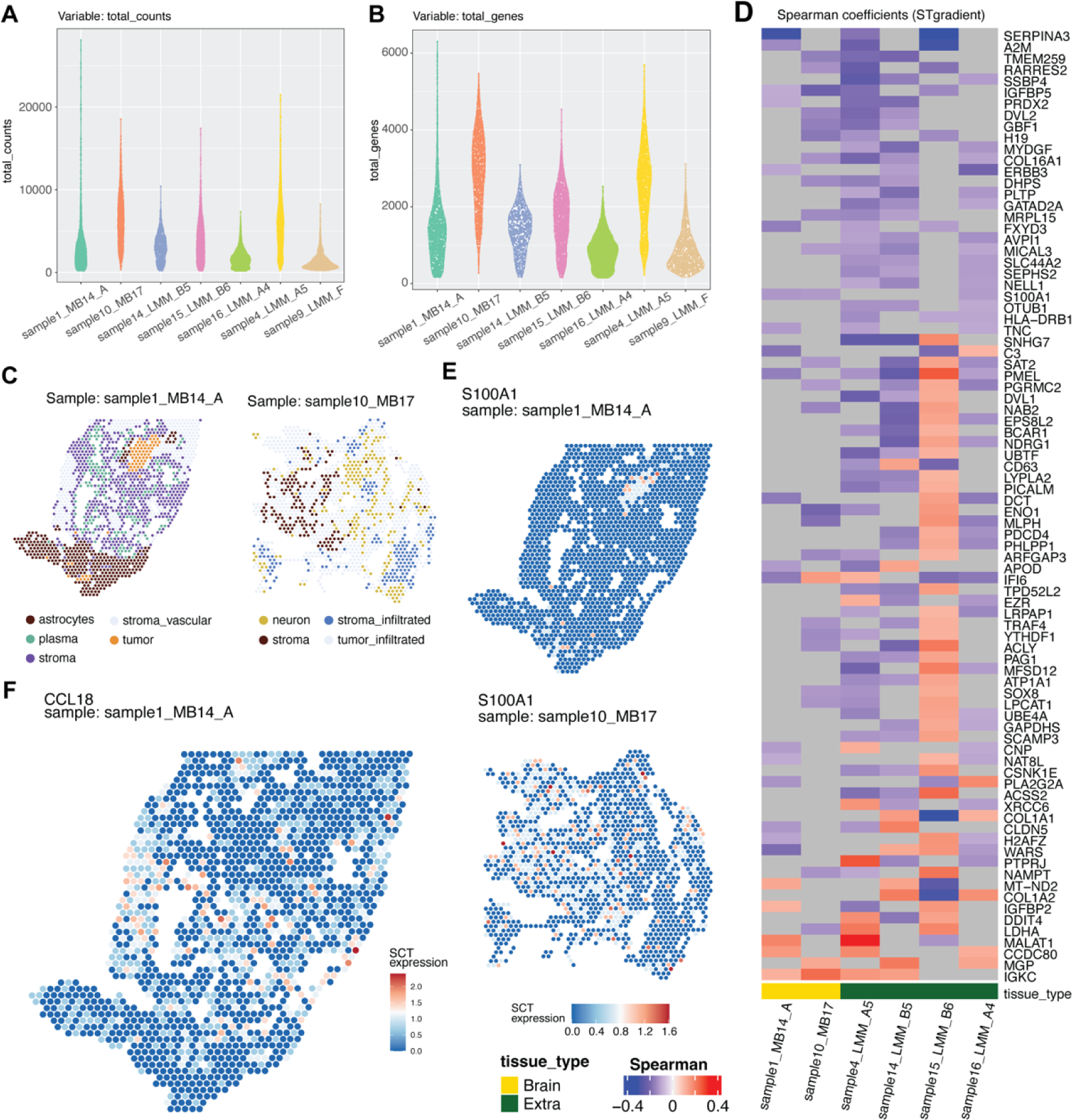
Analysis of LMM samples using spatialGE. Total number of counts per spot (A); Total number of non-zero genes per spot (B); Tissue domain classification classifications per spot based on STclust followed by differential expression analysis (C); Genes showing significant spatial gradients (STgradient, p-value < 0.05) and Spearman’s r > 0.1 (D). Expression of *S100A1* in two samples (E); Expression of *CCL18* in two samples (F). All figures (except panel D) were generated with spatialGE.

#### Merkel cell carcinoma

Merkel cell carcinoma (MCC) is a neuroendocrine skin cancer occurring mostly in patients of advanced age. Occurrence of MCC is rare, with survival rates depending mostly on the population or sex of the patients^22^. Recently, ST has been used to explore the tissue microenvironment of this understudied cancer type^23^. We obtained publicly available Merkel cell carcinoma data from lesions profiled with NanoString’s CosMx SMI technology. The data set comprised 95 FOVs from 11 tissue samples from 10 patients, allocated in six CosMx slides. We used spatialGE’s spatial analysis methods to investigate the differences in spatial expression in the context of response to immune checkpoint inhibitor therapy (administered after tissue sampling). The emergence of MCC tumors is associated with repeated exposure to ultraviolet light or infection with polyomavirus (MCPyV) ^22^. Hence, we also explored associations of spatial gene expression patterns with the MCPyV infection status.

The filtering procedure resulted in an average of 75 to 353 counts per cell across all 95 FOVs (**Fig. 4A**). Pseudo-bulk analysis (*QC and data transformation* module) revealed that gene expression of FOVs varied according to the level of response to immunotherapy (**Fig. 4B**). However, this pattern also overlaps partially with the differences observed among slides, highlighting potential batch effects (**Fig. 4B**). Identification of cells by Insitutype (*Phenotyping* module) identified as “Melanocytes.2” cells expressing keratin genes and *CHGA* (**Fig. 4C**). Hence, we reassigned those as cells as Merkel cells. These classifications serve as a basis for collapsing cell types into tumor (Merkel cells) or non-tumor regions. Next, we used the *Spatial gene set enrichment* module to test for the tendency of the expression of gene sets to form spatial “hotspots”. After summarizing FOVs to the tissue sample level, we observed that two out of four tissue slices from tumors that did not respond to immunotherapy showed spatial uniformity (i.e., non-significant STenrich test) in the expression of genes from the fatty acid metabolism (**Fig. 4D**). A negative MCPyV tissue sample from a tumor that responded to immunotherapy showed spatial uniformity in fatty acid metabolism gene expression (**Fig. 4D**).

**Figure 4.**
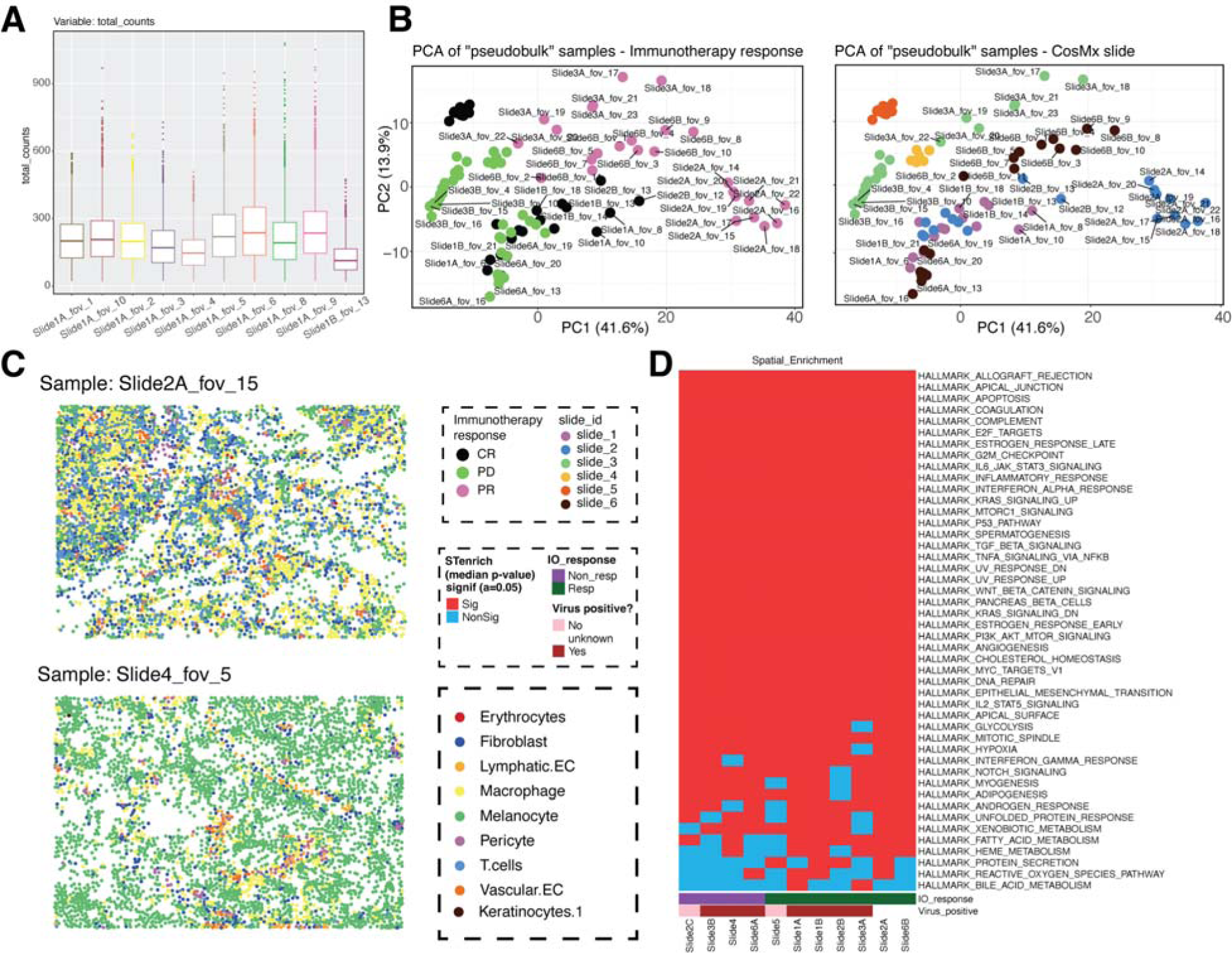
Analysis of Merkel cell carcinoma samples using spatialGE. Total number of counts per cell for 10 of the FOVs in the data set (A); A principal components plot resulting from pseudobulk analysis (B); Cell type classifications resulting from Insitutype (C); Hallmark gene sets showing evidence of spatially-restricted enrichment (D). The median p-value is presented for all FOVs within a tissue sample (STenrich, p-value < 0.05 = red, p-value >= 0.05 = blue). All figures (except panel D) were generated with spatialGE.

## Discussion

Given the rising popularity of ST in tissue biology research, especially in areas such as oncology and neurobiology^2, 24^, the demand for accessible and standardized analysis pipelines is also increasing. Nonetheless, the high dimensionality and spatial nature of these data sets require specialized help from bioinformaticians. The user-friendly interface and diverse modules in spatialGE cater to researchers with varying levels of programming skills, thereby democratizing access to exploratory data analysis involving ST experiments. With spatialGE, researchers with knowledge of tissue biology are now able to explore ST data and formulate hypotheses that can be translated into functional experiments. This will also allow efficient collaboration with bioinformaticians, who can be consulted for advanced and customized data analysis pipelines to follow up the spatial biology insights found from exploratory data analysis using spatialGE.

We applied spatialGE to analyze brain and extra-cranial melanoma metastases profiled with the 10X Genomics’ Visium technology. The visualizations in the *QC and data transformation* module were useful in identifying exceptionally lower gene counts in one of the brain metastasis samples (sample9_LMM_F), and hence this sample was removed from the spatial gradient analysis. The analysis also indicated that in the two brain melanoma metastases, the gene *S100A1* was expressed higher in spots near the tumor areas. The *S100A1* is a member of a large family implicated in the proliferation of several types of cancer, including melanoma^25^. This finding also supports the idea that surrounding stromal elements in the tumor microenvironment in the central nervous system may support the proliferation of cancer cells, as noted in leptomeningeal metastatic melanoma^4^. We also observed marked spatial expression patterns in sample1_MB14_A, where a region of astrocytes was identified. The astrocyte region expressed *CHI3L1*, a chitinase involved in inflammation and tissue-repairing processes^26^. It has been found that GFAP+/pSTAT3+ astrocytes mark a subpopulation of reactive astrocytes which initiates a transcriptional program driving the secretion of *CHI3L1*, consequently leading to activation of the PI3K/AKT signaling cascade in metastatic cancer cells^27^. Astrocytes in the brain tumor microenvironment have been shown to form tumor-reactive glia in response to metastatic brain tumors^28, 29^. Especially, reactive astrocytes have been shown to confer a growth promoting and immunomodulatory function in the brain tumor microenvironment^30, 31^. The astrocyte-dominated regions not only expressed *CHI3L1* but, interestingly, *CHI3L2* expression was also revealed, suggesting that phenotypically similar RAs may be captured in this sample. We explored expression gradients associated with this astrocyte area and noticed the expression of the chemokine ligand *CCL18* increased with distance from this area. It has been hypothesized that *CCL18* acts as an immunosuppressor in tumors located in the brain^32^. However, our results indicating lower expression adjacent to astrocytes require further research.

We also illustrated the use of spatialGE to uncover patterns from a MCC study where data was generated using NanoString’s CosMx SMI platform. In the MCC study, the use of principal component analysis in the QC and data transformation module uncovered a potentially confounding interaction between the response to immunotherapy (administered after sample collection) and the CosMx slide that originated the data (i.e., technical effect). While batch effects are difficult to remove in ST data^6^, spatialGE assists users in revealing variables that could interfere with or impact downstream statistical analyses. We also noticed that expression of fatty acid metabolism genes was spatially aggregated in most MCC samples, except in two tissues from tumors non-responsive to immunotherapy and one immunotherapy-responsive tumor also positive to MCPyV. It has been suggested that MCC relies on fatty acid production for proliferation^33^. However, these results suggest that not only expression but uniform expression across the tumor may reduce the chances of immunotherapy success.

One of the major strengths of spatialGE is its robust web application. The use of dockerization allows each process to run in its own self-contained environment. This also facilitates the future integration of spatialGE with other analytical methods built in other languages, such as Python, C++, or Julia. This dockerized approach opens possibilities to allow integration with code from members of the research community. The application also includes features like email notification and the ability to save results in a project, enhancing the user experience. To further democratize the analysis of ST data, ongoing efforts are integrating spatialGE methods in other user-friendly tools such as WebMeV^34^ and GenePatten^35^. Lastly, future work is ongoing to incorporate additional analysis modules and functionality into spatialGE.

### Conclusions

With the increasing popularity of ST, challenges have emerged regarding the analysis of this multimodal omics technology. Not only does ST analysis require methods adjusted to the spatial properties of the data (e.g., spatial autocorrelation), but also the need for standardized and accessible pipelines that are readily available to the entire research community and not only to scientists with programming skills. We have presented a solution to these challenges by providing a point-and-click, user-friendly web application to allow all researchers to conduct spatial analysis, thus democratizing the analysis of ST data. The spatialGE web application facilitates comparative analysis among multiple samples and can take metadata (e.g., clinical outcomes), enabling hypothesis generation. We have shown the utility of spatialGE to develop novel biological hypotheses through the analysis ST data from two cutaneous cancers studies. These hypotheses could then be further explored with additional experiments and more advanced bioinformatic analyses.

## Methods

spatialGE is a “point-and-click” wrapper interacting with the spatialGE R package^12, 36^. We have also expanded its functionality by implementing SpaGCN^15^ for tissue domain detection, STdeconvolve^17^ for reference-free deconvolution, and Insitutype^19^ for cell-level phenotyping. spatialGE was developed in a modular framework, intending to guide users through a ST data analysis workflow. The analytical workflow starts with an intuitive data input interface, which enables the data filtering, quality control, and normalization module. Once the data sets are filtered and/or normalized, modules for spatial heterogeneity quantification, tissue domain identification, cell phenotyping, and spatial gene set enrichment are enabled. Completion of domain identification will enable differential gene expression and gradient analyses (**Fig. 2**). Data uploaded by a user is kept within a project that is held within a password-protected account.

### Software architecture

spatialGE is currently hosted on an external facing Moffitt Cancer Center server with public access. The server runs Apache (https://httpd.apache.org/), PHP (https://www.php.net/), and MySQL-server (https://www.mysql.com/), providing support for the Laravel framework (https://laravel.com/) used in the development of the backend to handle user requests (e.g., text inputs, analysis requests) and store data such as user accounts, project information, and metadata associated with the spatial transcriptomics. The spatialGE application has been developed with portability in mind, achieved via containerization with Docker (https://www.docker.com/). When an analysis request is started in the web interface, the Docker container is initialized and runs the spatialGE R package (or other modules incorporated in the web interface, like SpaGCN Python package) and other libraries necessary to complete the analysis (**Fig. 5**). Analyses are queued and processed using Supervisor (http://supervisord.org/), allowing to better use and manage physical resources like RAM and CPUs. User parameters are captured through the web interface for each analysis and then fed into an R/Python template, the resulting script is executed inside a container with access to data using a mounted volume, this allows the user to access the results in the web application once completed, making spatialGE an interface to ease the interaction with R/Python packages that would otherwise require technical skills. Users can opt to receive email notifications when jobs are completed, allowing the web browser to be closed. The email notification contains a link to the project, with results displayed in the corresponding module.

**Figure 5.**
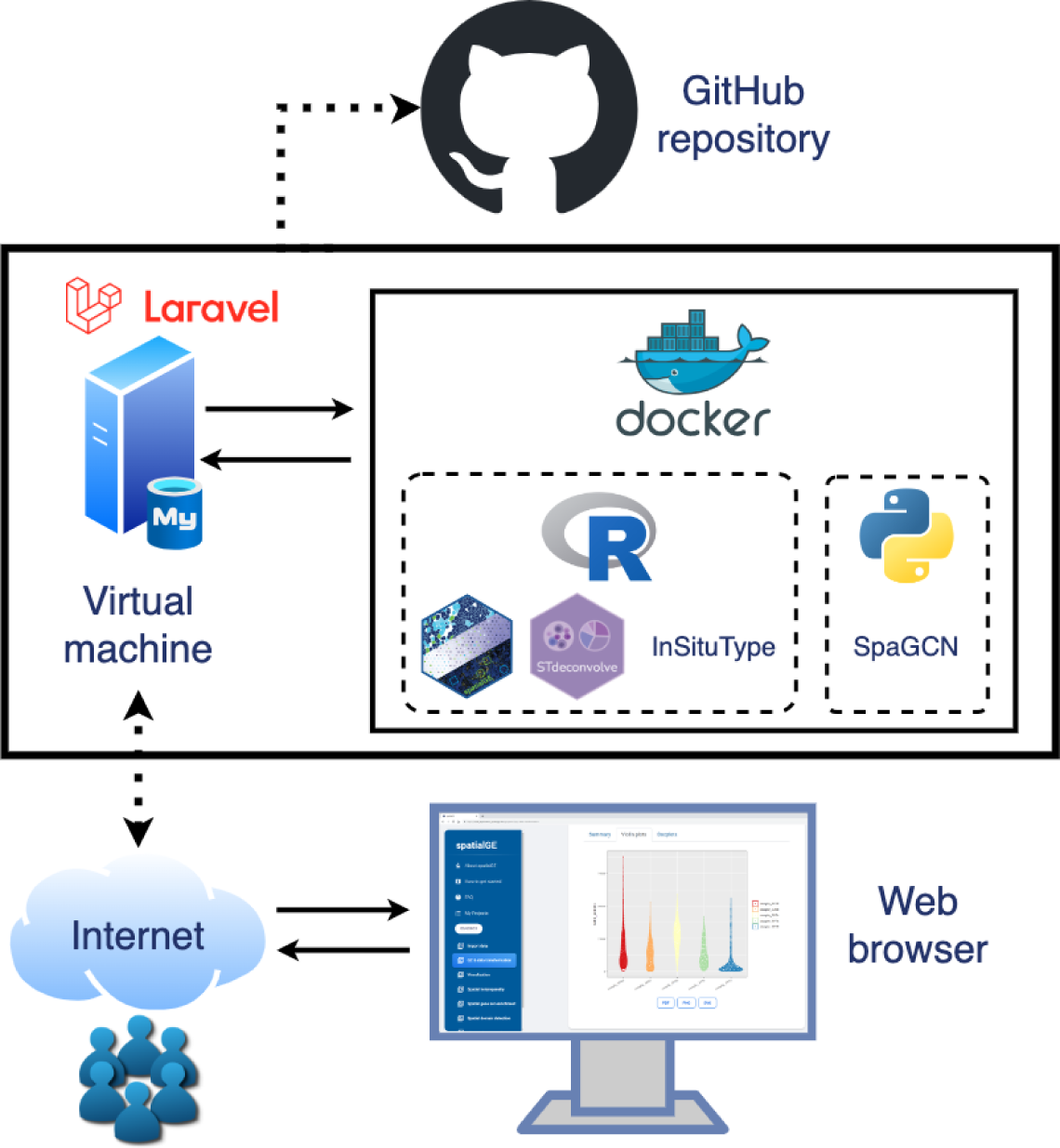
A diagram of the architecture of the spatialGE web application. Analysis requests are sent by the users through the web browser to the server, where a Docker container runs either R or Python scripts to generate the results, which are then accessible via the web browser by users.

The spatialGE R package run in the back end of the web application is available as a GitHub repository (see Availability of data and materials) and can also be run as a standalone library from the R console. The STlist S4 data structure stores raw gene expression counts and spatial coordinates. Most functions of the spatialGE R package return an updated STlist containing the results of the statistical analysis performed. Functions for normalization and log-transformation, visualization (“quilt” plots and expression surfaces), spatial heterogeneity statistics (Moran’s I/Geary’s C), and tissue domain detection (STclust) were presented in the original release of spatialGE. As part of the release presented in this work, the spatialGE R package now includes functions for the detection of gene expression spatial gradients (STgradient), spatial gene set enrichment (STenrich), differential expression (STdiff) (see Methods – *spatialGE analysis modules*).

### spatialGE analysis modules

spatialGE and the accompanying spatialGE R package feature three novel methods for studying spatial patterns in ST samples (STdiff, STgradient, and STenrich). The *Differential expression* module (STdiff^16^ function) allows testing for differentially expressed genes among tissue domains after running the *Spatial domain detection* or *Phenotyping* modules. The *Differential expression* module fits linear mixed models with an exponential covariance structure to test for differences in gene expression among the tissue domains or cell phenotypes. A non-spatial alternative that fits traditional Wilcoxon’s or t-tests is also provided. spatialGE also features integration with the Python package SpaGCN^15^ for tissue domain detection, which allows users to execute the *Differential expression* module on tissue domains inferred by SpaGCN.

The *Spatial gradients* module (STgradient function; **Additional file 4A**) calculates Spearman correlation coefficients between gene expression and the average or minimum distance of each spot/cell to a reference tissue domain. The *Spatial gradients* module allows for the specification of a minimum number of neighboring spots/cells belonging to the reference domain for the spot/cell to be considered in the calculation of distances (**Additional file 4A**). Given the excess of zero counts in spatial transcriptomics data, users can opt to exclude outlier spots/cells from the calculations or use robust regression as implemented via the MASS^37^ and sfsmisc^38^ packages.

The *Spatial gene set enrichment* module (STenrich function; **Additional file 4B**) allows users to test for the occurrence of “hot spots” in the expression of a gene set within each sample using a modified approach presented in a previous study^39^. For a given gene set, the average gene set expression is calculated for each spot/cell, and those spots/cells with expression or score above a user-specified threshold (average expression plus 1.5 standard deviations) are identified, and the sum of the spatial Euclidean distances between these identified spots/cells is calculated. Then, an equal number of spots/cells are selected from the whole sample, and the sum of their spatial Euclidean distances is calculated. The random selection is repeated a number of times (e.g., 1000 times), generating a null distribution to test if the sum of distances between the high expression spots is significantly smaller than the sum of distances among the randomly selected spots (**Additional file 4B**).

In addition to STdiff, STgradient, and STenrich, spatialGE supports the other processing and analysis methods in the spatialGE R package^12^. The *QC & data transformation* module allows the removal of genes or spots/cells for each sample using an interface with sliders to set minimum and maximum thresholds. The module supports the generation of principal component plots of “pseudobulk” counts^12^ to assess overall transcriptomic similarities among the samples. Currently, spatialGE supports normalization via library size scaling followed by log transformation and SCTransform^13^. In the *Visualization* module, the users can generate plots of gene expression via “quilt” plots showing raw or transformed gene expression values at each spot/cell or gene expression surface plots using “kriging” or spatial-interpolation^12^. The *Spatial heterogeneity* module calculates global Moran’s I and Geary’s C for a series of user-specified genes and across samples in a project and creates a plot to qualitatively detect patterns of associations between the existence of spatial patterns in gene expression and sample-level variables (e.g., tissue type, treatment). To complement spatialGE’s functionality, SpaGCN has been integrated into the platform, allowing users to choose between this graph convolutional network method and spatialGE’s native solution (STclust). Biological identification of spots or cells can be achieved with STdeconvolve (Visium) or InSituType (CosMx). Several cell type signatures are built, so that the user can select the most appropriate for the analyzed tissue type.

### Software usability and reproducibility testing

#### User Advisory Board

Throughout the development of spatialGE, feedback on the usability and utility of the application by members of the scientific community with a diverse array of skills and whose research involves analysis of spatial tissue biology. The spatialGE User Advisory Board (UAB) includes tissue biologists, pathologists, basic scientists, clinicians, and bioinformaticians, with varying computer programming skills. The UAB provided feedback to the spatialGE implementation team during group and individual meetings. This feedback was critical for developing user-friendly software that non-data scientists would want to use for their ST studies.

#### Robustness and reproducibility

We tested the stability and robustness of spatialGE by simulating a high usage period. A Python framework was developed to simulate eight users running jobs simultaneously within spatialGE. The workflow used Selenium WebDriver (https://www.selenium.dev/) to simulate user actions such as clicking buttons, scrolling, selecting files for upload, and any additional actions a user carries when using the spatialGE web app. We simulated three scenarios based on the memory usage of the requested job: 1) Four users utilizing the *Visualization* module (“light” job), two users normalizing data (“heavy” job), and two users utilizing the *Spatial heterogeneity* module (“medium” job); 2) two users in the *Visualization* module, four normalizing data, and two in *Spatial heterogeneity*; and 3) two users in the *Visualization* module, two normalizing data, and four in *Spatial heterogeneity*. The simulations showed that our process queuing system allocated resources with minimal wait times, except in one normalization and one visualization case (**Additional file 5A**). The time to process was consistent for each process, with the most variation in normalization (**Additional file 5B**). Memory usage was also variable in normalization and the *Spatial heterogeneity* module (**Additional file 5C**).

Another key aspect during the development of scientific software is ensuring the reproducibility of analytical tasks. To this end, spatialGE has built-in functionality to retrieve the parameters used to execute the analyses, enabling users to re-run jobs if needed and replicate their results. For each module within the web application, users can download tables and figures in raster and vector formats (i.e., PNG, PDF, and SVG).

### Melanoma metastasis study

Frozen tissue specimens from melanoma brain metastases were processed according to the Visium Spatial Gene Expression User Guide using reagents from the Visium Spatial Gene Expression Kit (10X Genomics, Pleasanton, CA). Three tissue slices from three melanoma brain metastasis sites were processed. For comparative analysis, we also included data from four extra-cranial metastatic melanoma tissue slices previously generated^4^ (GEO accession: GSE250636). Our study followed recognized ethical guidelines (e.g., Declaration of Helsinki, CIOMS, Belmont Report, U.S. Common Rule). The melanoma brain metastasis samples were procured during routine surgical resection and post-mortem under protocols approved by the Institutional Review Board (MCC 21044 and MCC 20779).

After sequencing on two separate Illumina NextSeq 2000 runs, the .fastq files were created using the default settings in Space Ranger mkfastq from the combined base call intensities outputs from the two runs. The reads were processed and aligned to the h38 human genome with *space ranger count*. The resulting count matrices contain the number of RNA counts for each gene at each spot, along with the associated spot coordinates. The data from the three melanoma brain metastatic specimens collected for this study will be made publicly available at the time of publication. A list of all the samples and accession numbers is in the **Additional file 6**. The .h5 files, spot coordinates, and images were uploaded into spatialGE. We have made available the parameter files provided by the application at each step of the analyses, which are available as Supplementary material for reproducibility within spatialGE (see **Additional file 7**).

First, we performed spot and gene filtering and normalization (SCTransform). Next, we applied STclust to define tissue niches and differential expression analysis (Wilcoxon’s rank sum test) to find genes highly expressed in each tissue domain. We renamed the domain(s) with high expression of melanoma genes (*MLANA*, *PMEL*, *TYR*, *TYRP1*) as “tumor”. Otherwise, the domain was re-labeled as “non-tumor”. Lastly, we tested for spatial gene expression gradients (STgradient) using the minimum distance to tumor and tested for spatial gene set enrichment (STenrich) across each entire tissue slice.

### Merkel cell carcinoma study

We used spatialGE to analyze Merkel Cell carcinoma (MCC) data generated with the CosMx platform. The MCC data set contained 95 fields of view (FOVs) originating from 11 tissue slices from 10 patients arranged in six slides. The fresh tumor tissue samples were collected from patients with Merkel cell carcinoma at the H. Lee Moffitt Cancer Center following written informed consent under the IRB-approved Total Cancer Care Protocol (MCC 14690, 94962), in accordance with the Declaration of Helsinki. The data will be publicly available at the time of publication, and a list of accession numbers and associated metadata can be found in **Additional file 6**.

For each CosMx slide, we uploaded the gene expression files (*_exprMat_file.csv), and cell coordinates (*_metadata_file.csv) into spatialGE. We have made available the parameter files provided by the application at each step of the analyses, which are available as Supplementary material for reproducibility within spatialGE (see **Additional file 7**). First, we performed cell and gene filtering and normalization (SCTransform). Next, we defined cell types using the InSituType^19^ implementation available in spatialGE. After differential expression analysis, we noticed that the label “Melanocytes.2” showed high expression of *CHGA*, a marker of Merkel cells. Hence, we renamed “Melanocytes.2” as “tumor”. Lastly, we tested for spatial gene set enrichment (STenrich) using spatialGE.

## Additional files

### Additional file 1

Compressed cell type signatures used for deconvolution of Visium data in spatialGE with STdeconvolve.

### Additional file 2

Spatial piechart presenting the estimated proportion of cell types to each spot in sample1_MB14_A. Estimation was performed in the *Phenotyping* module with STdeconvolve. **Additional file 3:**

Compressed file containing Excel workbooks with the *Spatial gradients* (STgradient) module results for all melanoma brain metastasis (Visium) samples. An additional workbook is included with the results testing from gradients of expression with respect to the astrocyte-dominated region in sample1_MB14_A.

### Additional file 4

Diagrams of two new algorithms to test for spatial gene expression patterns in ST data. A) ST gradient. B) STenrich.

### Additional file 5

The wait time (A), time to complete a process (B), and maximum used RAM (C) during the simulations performed to test the spatialGE server’s robustness.

### Additional file 6

Excel workbook containing the sample metadata used in spatialGE analyses. The file also contains the accession numbers and links to the data used in this work.

### Additional file 7

Files output by spatialGE containing the parameters used during the execution of the modules to analyze the data sets.

## Acknowledgments

This research was supported by the Biostatistics and Bioinformatics Shared Resource at the H. Lee Moffitt Cancer Center & Research Institute, an NCI-designated Comprehensive Cancer

Center (P30 CA076292). We thank Dr. Mingyao Li (University of Pennsylvania) for support in integrating SpaGCN in spatialGE.

## Author contributions

### Funding

National Institutes of Health – NIH (U01 CA274489, T32 CA23339).

### Availability of data and materials

The extracranial metastatic melanoma Visium samples are available at the Gene Expression Omnibus (GEO accession GSE250636). The brain metastatic melanoma (Visium) and Merkel cell carcinoma (CosMx) data will be available at time of publication. The spatialGE web application is available at spatialGE.moffitt.org. The source code is stored as a GitHub repository (https://github.com/FridleyLab/web_application_spatialGE). The spatialGE R package running in the back end of the application can be installed for command-line use from https://github.com/FridleyLab/spatialGE.

## Ethics declarations

### Ethics approval and consent to participate

Our study followed recognized ethical guidelines (e.g., Declaration of Helsinki, CIOMS, Belmont Report, U.S. Common Rule). Sample collection followed protocols approved by the Institutional Review Board at Moffitt Cancer Center (MCC 21044, MCC 20779, MCC 14690, and MCC 94962).

## Figures

**Additional file 2.**
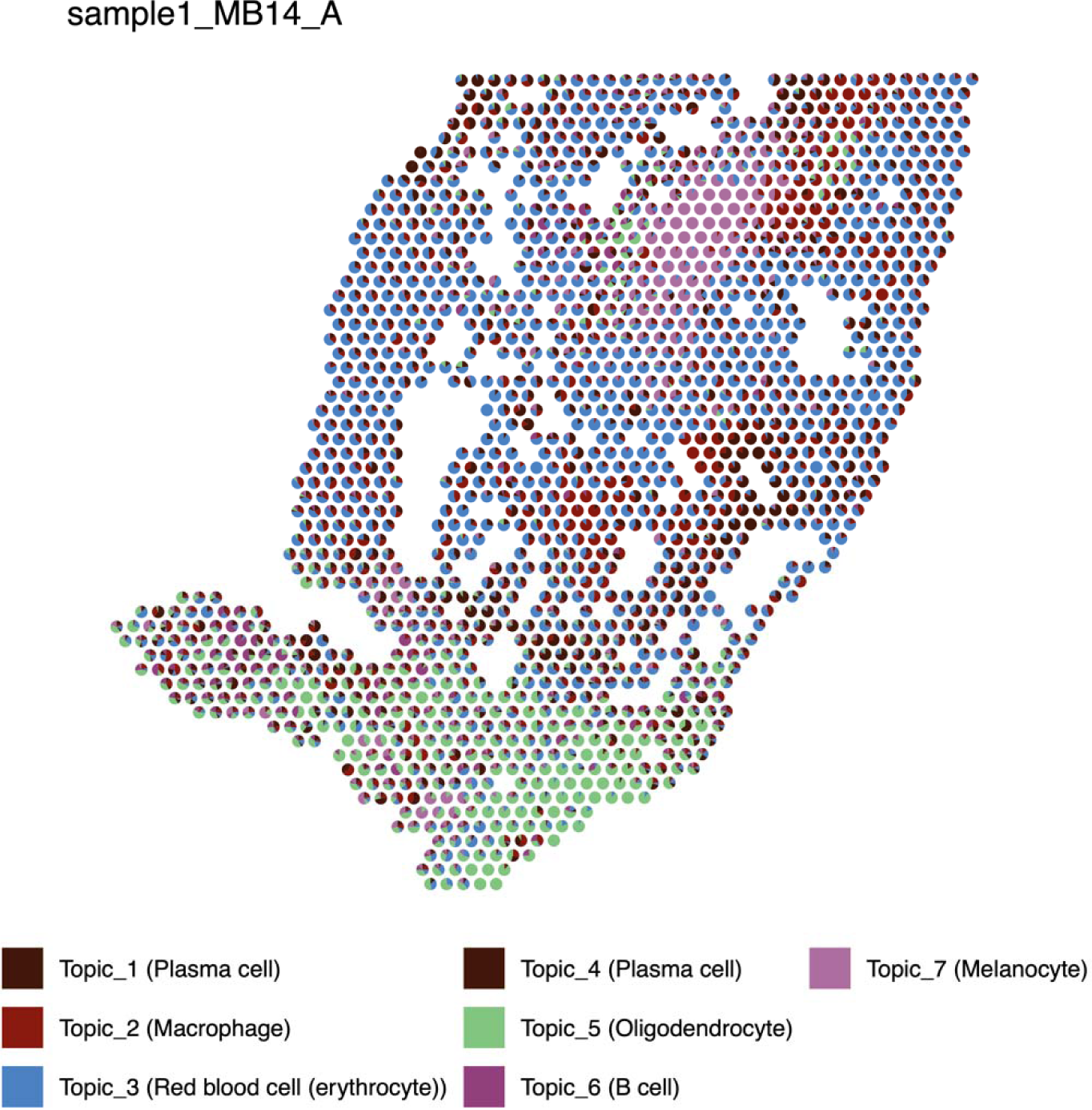
Spatial piechart presenting the estimated proportion of cell types to each spot in sample1_MB14_A. Estimation was performed in the *Phenotyping* module with STdeconvolve.

**Additional file 4.**
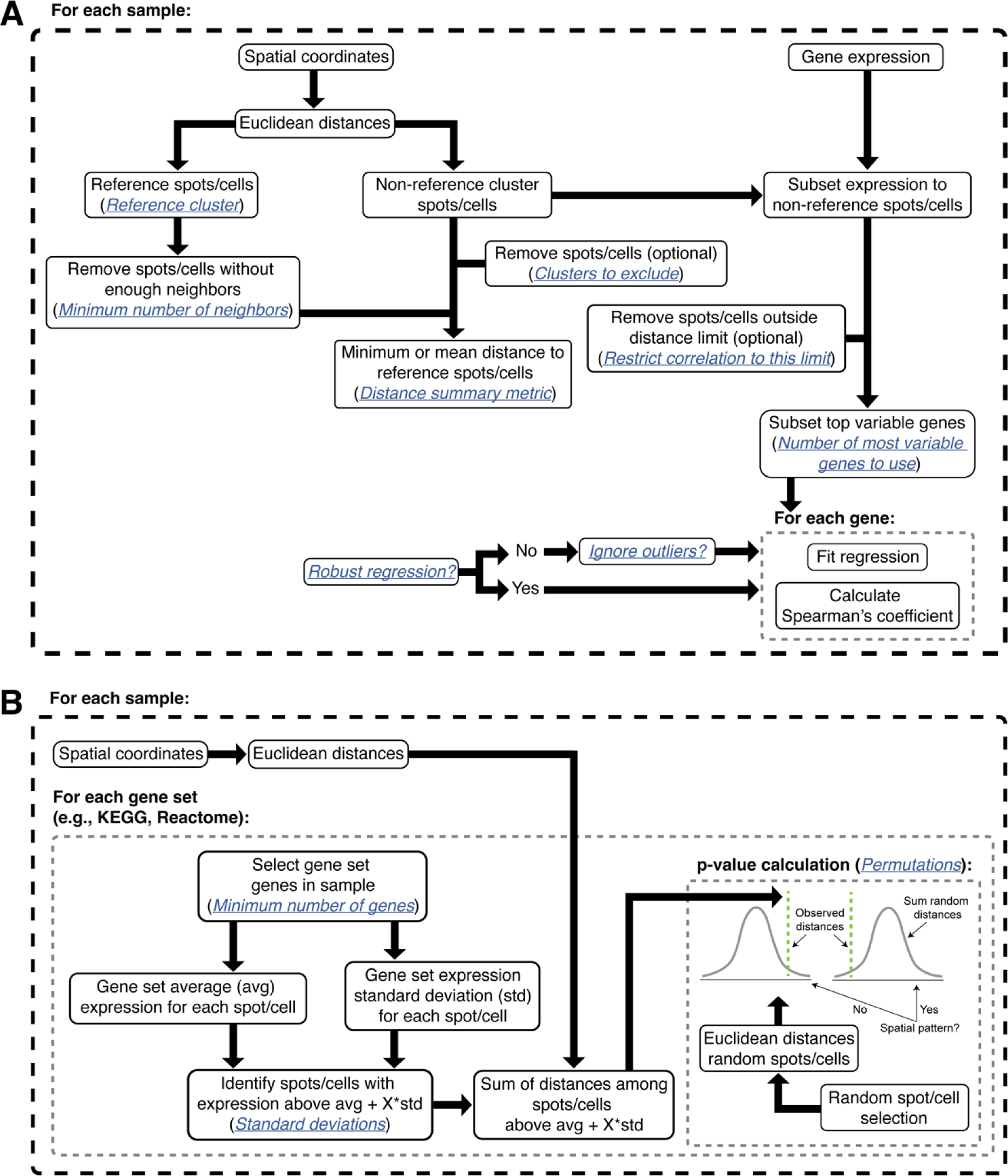
Diagrams of two new algorithms to test for spatial gene expression patterns in ST data. A) ST gradient. B) STenrich.

**Additional file 5.**
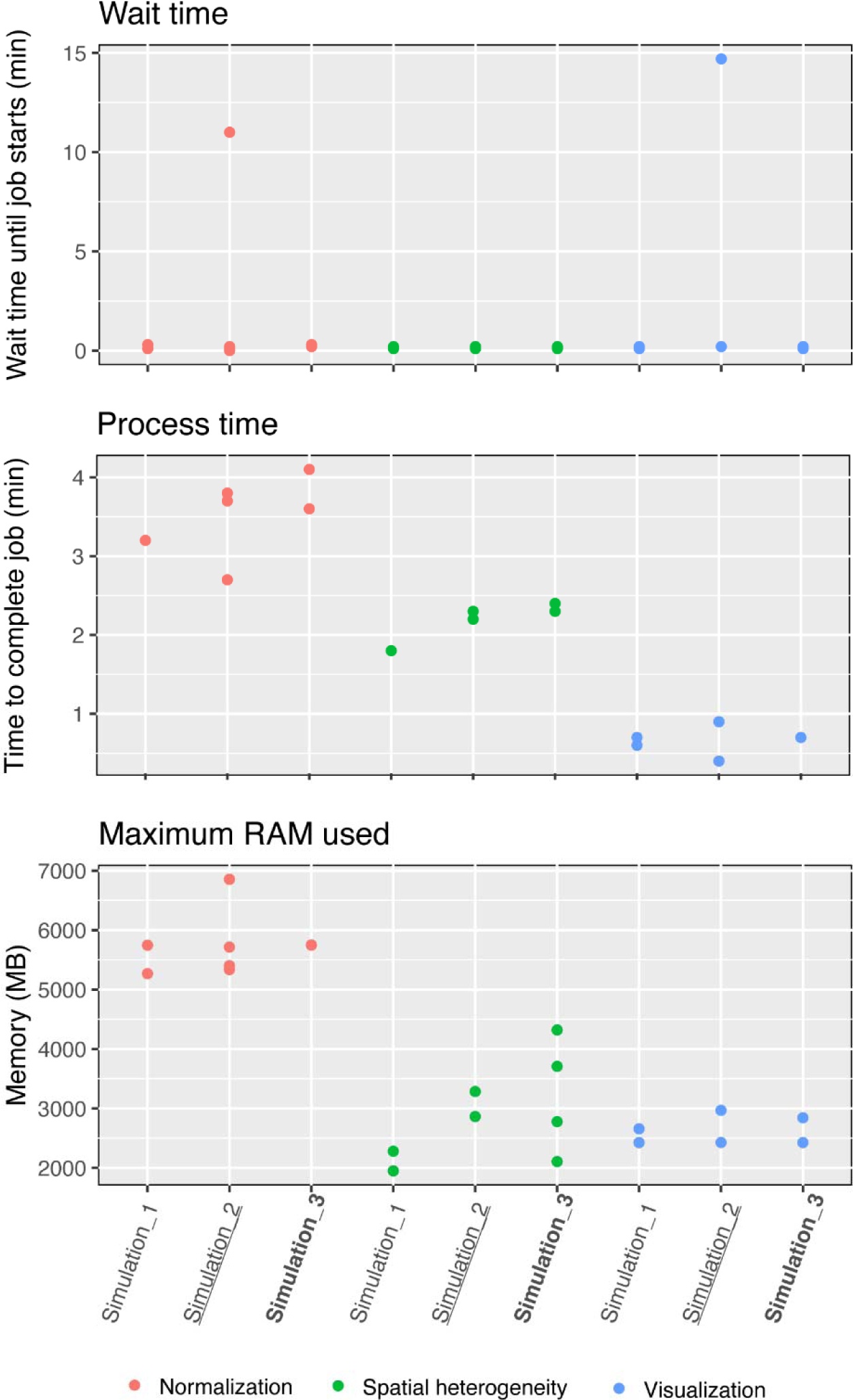
The wait time (A), time to complete a process (B), and maximum used RAM (C) during the simulations performed to test the spatialGE server’s robustness.

## References

1. Wang Y, et al. Spatial transcriptomics: Technologies, applications and experimental considerations. Genomics 115, 110671 (2023).

2. Moses L, Pachter L. Museum of spatial transcriptomics. Nat Methods 19, 534–546 (2022).

3. Rao A, Barkley D, Franca GS, Yanai I. Exploring tissue architecture using spatial transcriptomics. Nature 596, 211–220 (2021).

4. Alhaddad H, et al. Spatial transcriptomics analysis identifies a tumor-promoting function of the meningeal stroma in melanoma leptomeningeal disease. Cell Rep Med, 101606 (2024).

5. Wang Q, et al. Spatially Resolved Transcriptomics Technology Facilitates Cancer Research. Adv Sci (Weinh) 10, e2302558 (2023).

6. Fang S, et al. Computational approaches and challenges in spatial transcriptomics. Genomics Proteomics Bioinformatics, (2022).

7. Liu B, Li Y, Zhang L. Analysis and Visualization of Spatial Transcriptomic Data. Front Genet 12, 785290 (2021).

8. Keller MS, Gold I, McCallum C, Manz T, Kharchenko PV, Gehlenborg N. Vitessce: integrative visualization of multimodal and spatially-resolved single-cell data.

9. Dries R, et al. Giotto: A toolbox for integrative analysis and visualization of spatial expression data. Genome Biol 22, 78 (2021).

10. Yang S, Zhou X. SRT-Server: powering the analysis of spatial transcriptomic data. Genome Med 16, 18 (2024).

11. Wang G, Wu S, Xiong Z, Qu H, Fang X, Bao Y. CROST: a comprehensive repository of spatial transcriptomics. Nucleic Acids Res 52, D882–D890 (2024).

12. Ospina OE, et al. spatialGE: Quantification and visualization of the tumor microenvironment heterogeneity using spatial transcriptomics. Bioinformatics 38, 2645– 2647 (2022).

13. Hafemeister C, Satija R. Normalization and variance stabilization of single-cell RNA-seq data using regularized negative binomial regression. Genome Biol 20, 296 (2019).

14. Cressie NAC. Statistics for spatial data. Wiley & Sons (1993).

15. Hu J, et al. SpaGCN: Integrating gene expression, spatial location and histology to identify spatial domains and spatially variable genes by graph convolutional network. Nat Methods 18, 1342–1351 (2021).

16. Ospina OE, Soupir AC, Manjarres-Betancur R, Gonzalez-Calderon G, Yu X, Fridley BL. Differential gene expression analysis of spatial transcriptomic experiments using spatial mixed models. Sci Rep 14, 10967 (2024).

17. Miller BF, Huang F, Atta L, Sahoo A, Fan J. Reference-free cell type deconvolution of multi-cellular pixel-resolution spatially resolved transcriptomics data. Nat Commun 13, 2339 (2022).

18. Hu C, et al. CellMarker 2.0: an updated database of manually curated cell markers in human/mouse and web tools based on scRNA-seq data. Nucleic Acids Research 51, D870–D876 (2023).

19. Danaher P, et al. Insitutype: likelihood-based cell typing for single cell spatial transcriptomics. bioRxiv, (2022).

20. Danaher P, et al. Advances in mixed cell deconvolution enable quantification of cell types in spatial transcriptomic data. Nat Commun 13, 385 (2022).

21. Rishi A, Yu HM. Current Treatment of Melanoma Brain Metastasis. Curr Treat Options Oncol 21, 45 (2020).

22. Becker JC, Stang A, Schrama D, Ugurel S. Merkel Cell Carcinoma: Integrating Epidemiology, Immunology, and Therapeutic Updates. Am J Clin Dermatol, (2024).

23. Zhang Y, et al. 297 Expanded genomic landscape of merkel cell carcinoma identifies new drivers. Journal of Investigative Dermatology 143, (2023).

24. Zhao SH, et al. A Bibliometric Analysis of the Spatial Transcriptomics Literature from 2006 to 2023. Cell Mol Neurobiol 44, 50 (2024).

25. Hua X, Zhang H, Jia J, Chen S, Sun Y, Zhu X. Roles of S100 family members in drug resistance in tumors: Status and prospects. Biomed Pharmacother 127, 110156 (2020).

26. Zhao T, Su Z, Li Y, Zhang X, You Q. Chitinase-3 like-protein-1 function and its role in diseases. Signal Transduct Target Ther 5, 201 (2020).

27. Dankner M, et al. Invasive growth of brain metastases is linked to CHI3L1 release from pSTAT3-positive astrocytes. Neuro Oncol 26, 1052–1066 (2024).

28. Fitzgerald DP, et al. Reactive glia are recruited by highly proliferative brain metastases of breast cancer and promote tumor cell colonization. Clin Exp Metastasis 25, 799–810 (2008).

29. Wingrove E, et al. Transcriptomic Hallmarks of Tumor Plasticity and Stromal Interactions in Brain Metastasis. Cell Rep 27, 1277–1292 e1277 (2019).

30. Qu F, et al. Crosstalk between small-cell lung cancer cells and astrocytes mimics brain development to promote brain metastasis. Nat Cell Biol 25, 1506–1519 (2023).

31. Wasilewski D, Priego N, Fustero-Torre C, Valiente M. Reactive Astrocytes in Brain Metastasis. Front Oncol 7, 298 (2017).

32. Henrik Heiland D, et al. Tumor-associated reactive astrocytes aid the evolution of immunosuppressive environment in glioblastoma. Nat Commun 10, 2541 (2019).

33. Landes JR, et al. Effect of selinexor on lipogenesis in virus-positive Merkel cell carcinoma cell lines. Clin Exp Dermatol 48, 903–908 (2023).

34. Wang YE, Kutnetsov L, Partensky A, Farid J, Quackenbush J. WebMeV: A Cloud Platform for Analyzing and Visualizing Cancer Genomic Data. Cancer Res 77, e11–e14 (2017).

35. Reich M, Liefeld T, Gould J, Lerner J, Tamayo P, Mesirov JP. GenePattern 2.0. Nat Genet 38, 500–501 (2006).

36. Ospina OE, Soupir AC, Fridley BL. spatialGE: An R package for visualization and analysis of spatially-resolved gene expression.). 1.2 edn (2023).

37. Venables WN, Ripley BD. *Modern applied statistics with S-PLUS*. Springer Science & Business Media (2013).

38. Maechler M, et al. Package ‘sfsmisc’. (2023).

39. Hunter MV, Moncada R, Weiss JM, Yanai I, White RM. Spatially resolved transcriptomics reveals the architecture of the tumor-microenvironment interface. Nat Commun 12, 6278 (2021).

